# High-frequency cortical neural inputs to muscles during movement cancellation

**DOI:** 10.1101/2023.08.09.552619

**Authors:** B Zicher, S Avrillon, J Ibáñez, D Farina

## Abstract

Cortical beta (13-30 Hz) and gamma (30-60 Hz) oscillations have been investigated during motor processing. Although they are at frequencies greater than the dynamic bandwidth of muscle contraction, these oscillations are partly transmitted from the cortex to motoneurons and muscles. Little is known about when and why this transmission occurs. We developed an experimental approach to examine these high frequency inputs to motoneurons under different motor states while maintaining a stable force, thus constraining behaviour. We acquired brain and muscle activity during a ‘GO’/’NO-GO’ task. In this experiment, the effector muscle for the task (tibialis anterior) was kept tonically active during the trials, while participants (N=9) reacted to sequences of auditory stimuli by either keeping the contraction unaltered (‘NO-GO’ trials), or by quickly performing a ballistic contraction (‘GO’ trials). Motor unit (MU) firing activity was extracted from high-density surface and intramuscular electromyographic signals, and the changes in its spectral contents in the ‘NO-GO’ trials were analysed. We observed an increase in beta and low-gamma (30-45 Hz) activity post ‘NO-GO’ cue at the brain and muscle levels. There was also an increase in the activity within 8-12 Hz, which was only observed at the muscle level. Overall, our results suggest that the cortical processing of movement cancellation occurs at least in part via increased power of high-frequency oscillations transmitted downstream to the muscles. These changes occur without alterations in behaviour, suggesting that the downstream transmission of these high-frequency oscillations does not have a direct functional impact.

## Introduction

Oscillatory synchronisation is a mechanism through which the activity of a population of neurons can be modulated. In the context of motor control, this mechanism has been previously reported in preparatory tasks (Schoffelen et al., 2005). Brain states associated to movement preparation (Kaufman et al., 2014; Schoffelen et al., 2005) allow us to investigate how neural communication is modulated between brain circuits and spinal outputs to muscles in conditions where the motor outputs do not change.

In human studies using magnetoencephalography (MEG) and electromyography (EMG), it has been shown that corticomuscular transmission in the gamma band (30-70 Hz) is increased during movement preparation (Schoffelen et al., 2005; Schoffelen et al., 2011). Beta oscillatory activity (13-30 Hz) has also been widely linked to sensorimotor processing, being prominent during isometric contractions (Baker, 2007; Engel and Fries, 2010; Kilavik et al., 2013). It has been further shown that beta oscillations are reliably transmitted to muscles, where, recently, they have been directly decoded from motor unit (MU) firing activity (Bräcklein et al., 2021; Bräcklein et al., 2022; Ibáñez et al., 2021). Furthermore, in the motor cortex, beta activity is reduced by movement initiation, and increased during movement cancellation (Barone and Rossiter, 2021; Kilavik et al., 2013; Wessel, 2020). Taken together, these results support the concept that high-frequency activity, which is outside the muscle bandwidth relevant for force control (Farina and Negro, 2015; Farina et al., 2014; Negro and Farina, 2011), presents prominent changes during various brain states associated to motor commands and modulates the communication between the brain and muscles.

While beta activity has been extensively investigated in the context of corticomuscular transmission during sustained contractions, less is known on neural connectivity between the brain and muscles during movement preparation and cancellation, i.e., when a movement is planned and then aborted. Cancelling prepared action plans is a particularly interesting motor state, since it is associated with inhibition of motor commands at the cortical level, thus transmission of high frequency oscillations to muscles does not seem functionally necessary.

Here, we study the modulation of motoneuron firing activity at the population level during periods of motor planning and action cancellation. Specifically, we designed a ‘GO’/’NO-GO’ paradigm in which the effector muscle was tonically active throughout the trials. Brain and muscle activity were concurrently recorded with electroencephalography (EEG) and EMG. The latter was decomposed into constituent spike trains using previously validated methods (Negro et al., 2016), to gain direct access to the MU firing activity at a population level. This allowed us to analyse the spectral contents of changes happening in the activity of pools of motoneurons (the neural drive to muscles) during sustained contractions, and the modulations of cortical rhythmic activity related to motor planning and action cancellation (in the case of ‘NO-GO’ trials). Additional computer simulations of a pool of motoneurons were also conducted to study how different types of neural inputs can modulate the motoneurons firing activity at the population level.

Overall, the study aimed to explore high-frequency modulatory inputs to motoneurons during changing brain states, that did not cause functional force modulations. The results provide, on one hand, new insights into the dynamic transmission of cortical inputs to spinal circuits, and, on the other hand, evidence for the possibility of detecting brain activity from peripheral interfaces.

## Materials and methods

### Computer simulations

Simulations of spinal motoneurons were run to explore how different low and high frequency inputs change their firing activity. The insights from the simulations were also useful when interpreting the results from the experimental part of this work.

A total of 30 S-type motoneurons were simulated to match the number of motor units identified during the experiments (see Results). Physiologically realistic two-compartmental models (soma and dendrite) were used, with parameters taken from a previously published work (Cisi and Kohn, 2008). The dendrite received synaptic input and leak currents, while the soma contained leak, Na^+^, fast K^+^ and slow K^+^ currents (Cisi and Kohn, 2008).

Common and independent inputs were generated as low-pass filtered Gaussian noises (common: < 3 Hz, independent: < 40 Hz).In scenarios where a specific oscillatory activity was transmitted to the pool, oscillations in the alpha, beta and gamma bands were represented as 300-ms long pure 10 Hz, 20 Hz and 35 Hz sinusoidal inputs with a constant amplitude that did not increase the discharge rate of motoneurons. A total of four scenarios were simulated, each with an event happening 3-s after the beginning of the simulation. The event occurs at this specific time to ensure that motoneurons were steadily firing. The first event consisted of a short drop in the low frequency common input. The other three events consisted of a burst of alpha, beta, or gamma oscillatory activity added to the common input, respectively. In all scenarios, the level of the independent input was chosen such that the coefficient of variation (CoV) of interspike intervals matched the experimental results (i.e., 13-16%). The amplitude of the total input was adjusted such that the average discharge rate of motoneurons matched the physiological values observed in the experimental part (i.e., 9-12 Hz).

### Experimental data acquisition

A total of 11 participants (all males, ages: 21-38 yrs) took part in this experiment and gave their written informed consent. Data from two participants were discarded from the analysis due to the presence of artifacts in the force signals. The study was conducted in accordance with the Declaration of Helsinki and was approved by the Imperial College Ethics committee (18IC4685).

#### Recordings

Experimental signals were non-invasively recorded from the brain using electroencephalography (EEG) and from the right tibialis anterior (TA) muscle using high-density surface EMG while participants performed isometric ankle dorsiflexions (Figure 1A). In two participants, multi-channel intramuscular EMG signals were also acquired.

**Figure 1:**
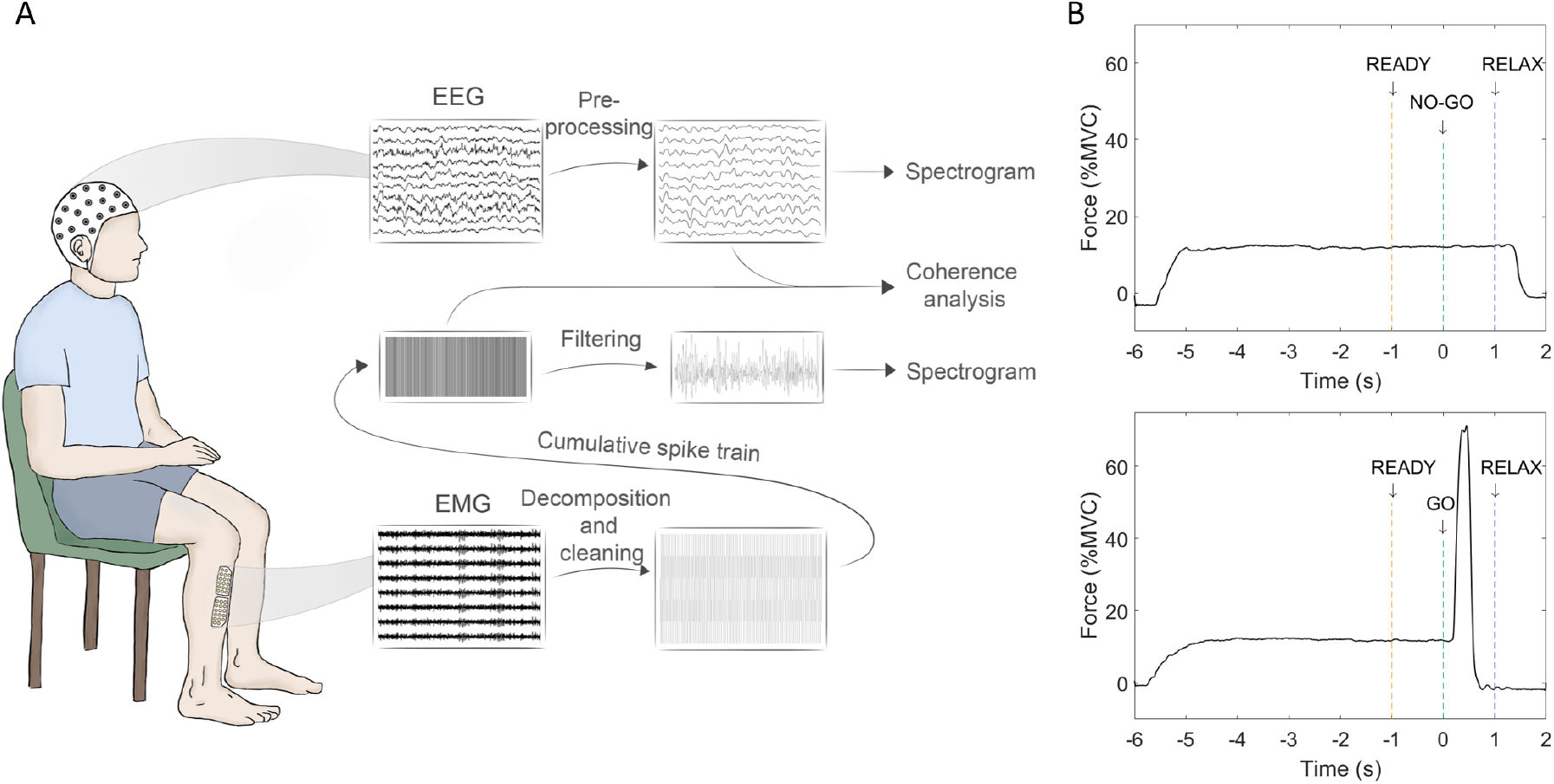
Schematic representation of the experimental protocol. A, EEG and EMG signals were concurrently recorded while participants were seated in a chair and were producing ankle dorsiflexions. EEG recordings were re-sampled and filtered offline (see Methods). EMG signals were decomposed into MU trains of action potentials, which were then summed to get the cumulative spike train (CST). Spectral and coherence analyses were performed on EEG and CST. B, Experimental tasks with ‘NO-GO’ (top figure) and ‘GO’ (bottom figure) trials. During each trial, the participants performed an isometric dorsiflexion at 10% MVC before reacting to an imperative stimulus. During ‘GO’ trials, participants performed a ballistic contraction, while during ‘NO-GO’ trials, they held the isometric contraction at 10% MVC until the last cue.

EEG signals were recorded using 31 active gel-based electrodes positioned according to the International 10–20 system with FCz used as reference (actiCAP, Brain Products GmbH), amplified and sampled at 1 kHz (BrainVision actiCHamp Plus, Brain Products GmbH). The signals were resampled at 2048 Hz offline.

Surface EMG was recorded using 64 channel grids (13×5 arrangement with one missing electrode in a corner) with an interelectrode distance of 4 mm (OT Bioelettronica). The signals were amplified, sampled at 2048 Hz (Quattrocento, OT Bioelettronica) and bandpass filtered (20-500 Hz). In seven participants, four surface grids were used to cover the TA muscle (Caillet et al., 2023). In the other two participants, two surface grids and respectively three and four intramuscular electrode arrays were used. These intramuscular arrays had 40 electrodes, with 20 electrodes located on each side of a flat thin-film and separated by an interelectrode distance of 500 μm (Muceli et al., 2022). In the experiments involving intramuscular recordings, all EMG signals were sampled at 10240 Hz, and intramuscular signals were bandpass filtered between 100 and 4400 Hz.

During the experiment, participants sat in a comfortable chair with their knee flexed at 75°, their right leg securely fixed to an ankle dynamometer with Velcro straps, and their foot positioned onto a pedal at 30° in the plantarflexion direction, 0° being the foot perpendicular to the shank. A force transducer (TF-022, CCT Transducer s.a.s) fixed to the pedal recorded the force. During all tasks, participants received visual feedback with a target representing the level of force to reach and a trace representing the force they produced.

EEG, EMG and force signals were synchronised using a common digital trigger signal sent to the systems.

#### Task

The task was based on a movement preparation and cancellation framework that allowed us to explore changes in spinal motoneuron population activity during these brain states.

At the beginning of the experiment, participants were asked to produce an ankle dorsiflexion contraction with maximal force to estimate their maximum voluntary contraction (MVC) force level. Then, the experiment was divided into three identical blocks with periods of rest in-between. Each block consisted of 35 trials, with one trial having a 50% chance of being a ‘GO’ or a ‘NO-GO’ trial. In both cases, throughout the trial, subjects received four auditory cues of various lengths and frequencies. Before the recordings started, subjects were familiarized with the task and practiced responding to the cues until they were confident in their understanding of the task. For each trial, a first cue indicated to the participants that they had to produce a 10% MVC force. Then, after 5 s, a second cue (warning stimulus) was given, and this was followed by a third cue (imperative stimulus) 1 s later, which informed the subjects about the trial type (‘GO’ or ‘NO-GO’). A fourth cue, 1 s after the imperative stimulus, indicated the end of trial. At this point participants had to relax and wait for the following trial (Figure 1B). During ‘GO’ trials, participants produced a ballistic contraction above 10% MVC as soon as possible after the imperative stimulus. During ‘NO-GO’ trials, participants kept the level of force at 10% MVC until the end of the trial indicated by the fourth cue. The timing of the cues was fixed during all trials to ensure that the evolution of the neural states decoded from motoneurons were temporally aligned across trials, thus allowing the averaging of the results. Participants only received visual feedback on the force produced until the third cue. Additional visual feedback on reaction time was presented to the participants after the ‘GO’ cues to ensure fast and consistent reaction times through each block. Trials were separated by a period of rest of 7 s on average with a variation of ±20%. One block of 35 trials lasted about 8 minutes.

### Data analysis

Spectral and coherence analyses were performed on signals recorded during ‘NO-GO’ trials, to be able to study the cancellation of a prepared movement. Force, EEG and EMG signals were cut -4 s to 1 s relative to the imperative cue before analysis. In addition, reaction time and CoV of force were analysed from ‘GO’ trials to check that the tasks were done correctly.

#### Force analysis

The force data was low-pass filtered with a cut-off frequency of 15 Hz using MATLAB *lowpass* function, baseline corrected, and normalised relative to the MVC. The reaction time was calculated during ‘GO’ trials as the duration between the imperative cue and the time point where the force signal exceeded 2 standard deviations of the mean force value in the 4-s interval preceding the ‘GO’ cue. The CoV of force was measured over a window of 4 s preceding the imperative cue. Only trials in which the force remained between 7% and 13% MVC in this interval were kept for further analysis.

To check the performance of the participants, the percentage of valid trials was quantified. A ‘GO’ trial was considered valid if the ballistic movement was ± 2 standard deviations of the participant’s mean reaction time. ‘NO-GO’ trials were considered valid if the force did not exceed 15% MVC throughout the trial. In addition, to discard from further analysis the trials in which participants did not perform the overall task optimally, specific criteria were defined. Trials were discarded if during the steady contraction part, force went 2% below or 2% above the average calculated across all trials, or if the force standard deviation was above 0.5% MVC. This ensured that only trials where participants kept the force at a relatively constant level were included in the analysis.

#### EMG decomposition

The EMG signals recorded from the different trials were concatenated and decomposed into constituent trains of action potentials using previously validated methods (Negro et al., 2016). The signals recorded by each grid were decomposed separately, and the discharge series of motor units identified in more than one grid were merged. The MU discharge times automatically detected by the algorithm were visually inspected and edited when necessary. The manual editing consisted of the removal of artifacts falsely identified as discharge times and the addition of discharge times missed by the automatic steps, followed by automatic verification of the validity of the edits (Hug et al., 2021).

#### Time-frequency analysis

The firing activity of simulated motoneurons were summed to estimate the cumulative spike train (CST). The CST was bandpass filtered between 8-45 Hz (3^rd^ order Butterworth) and its spectrogram was calculated using the MATLAB *spectrogram* function (rectangular window; segment length = 0.25 s; shift between adjacent segments = 10 samples).

When considering the experimental part, only the signals from the valid ‘NO-GO’ trials were included in the analysis (see the section “Force analysis”). Furthermore, MU spike trains from trials in which the COV of discharge rate was above 30% were also excluded.

For the simulations, the CST was calculated by summing the firing activity of the identified MUs. The CST was bandpass filtered depending on the bandwidth analysed (alpha: 8-12 Hz; beta: 13-30 Hz; gamma: 30-45 Hz) with a 3^rd^ order Butterworth filter.

EEG signals were first visually inspected, and trials in which movement artifacts were observed were excluded from the analysis, besides the ones that were not adequate according to the behavioural or MU discharge rate criteria (see above). The Laplacian derivation from channel ‘Cz’ was computed by subtracting the average electric potential recorded from the four closest equidistant channels, i.e., FC1, FC2, CP1, CP2 in our set-up. The surrogate ‘Cz’ channel was used to estimate the directional coherence. The resulting channel data was bandpass filtered before the analysis with a 3^rd^ order Butterworth filter according to the bandwidth under investigation.

For both signals (MU activity and EEG), the spectrograms were calculated with the segment length set to 0.25 s and a shift between adjacent segments of 10 samples. The results were averaged across trials before calculating the mean in each bandwidth (alpha: 8-12 Hz; beta: 13-30 Hz; gamma: 30-45 Hz). These values were then standardized in the window -3 s to 1 s relative to the ‘NO-GO’ cue. The average values for windows -2 s to -1 s, -1 s to 0 s and 0 s to 1 s were also calculated for each subject. To get the grand average results, first, for each subject separately, at each frequency sample, the average value was calculated in the window -3 s to -2 s. The value obtained was used as a reference of the baseline level of activity at each frequency examined. At each following time sample, the change relative to the baseline was calculated and the values were standardized. This represents the changes occurring relative to the baseline at each frequency for a subject. The mean of these results was then calculated to get the grand average.

#### Coherence analysis

The transmission of high frequency oscillations between the brain and muscles was studied by calculating the directional coherence between the ‘Cz’ channel and the CST. The EEG signal was processed as for the time-frequency analysis, except here a 1-45 Hz bandpass filter was used. The same trials were removed from the EEG and CST and then the signals were detrended. The coherence was calculated over 1-s windows (−2 s to -1 s; -1 s to 0 s; 0 s to 1 s relative to the imperative ‘NO-GO’ cue) using the Neurospec 2.11 toolbox coded for MATLAB (www.neurospec.org; Mathwoks Inc., USA). This toolbox uses the multitaper method (3 tapers) for the spectral estimation and allows for the estimation of directional coherence. The toolbox also calculates the 95% confidence limit for the coherence estimate that is used here to evaluate significance.

#### Statistics

The statistical analysis was performed with SPSS (IBM). Three main windows were compared: -2 s to -1 s (referred to as *baseline*), -1 s to 0 s (*preparation*) and 0 s to 1 s (*cancellation*) relative to the imperative ‘NO-GO’ cue. To test for significant changes in power in these time widows, repeated measures ANOVA was performed for each recording type and frequency band, with time as the within-subject factor. The same test was used to check for changes in average discharge rate. The sphericity was checked with the Mauchly’s test. Bonferroni correction was used to adjust for multiple comparisons. Results are reported as mean ± standard deviation, unless otherwise stated.

## Results

### Simulation results

Computer simulations were run to explore how the spiking activity of a population of motoneurons can be synchronized through various common inputs. The four scenarios simulated included one with a small decrease in the low frequency common input, and three cases with a short oscillatory input in alpha, beta and gamma, respectively.

Figure 2A on the left panel shows the common input to the pool of motoneurons in the first scenario, together with an example of the total input to one of the neurons. The event of a drop in input can be seen at second 0. The right panel in Figure 2A shows the spectrogram of the CST in the three bandwidths (alpha, beta, and gamma), calculated as the sum of the firing activity of the simulated motoneurons receiving the common input shown on the left panel. The spectrogram showed increased power around 11-12 Hz at the time of the drop in common input. This represents the synchronisation of the firing activity of multiple motoneurons at the frequency equal to their average discharge rate.

**Figure 2:**
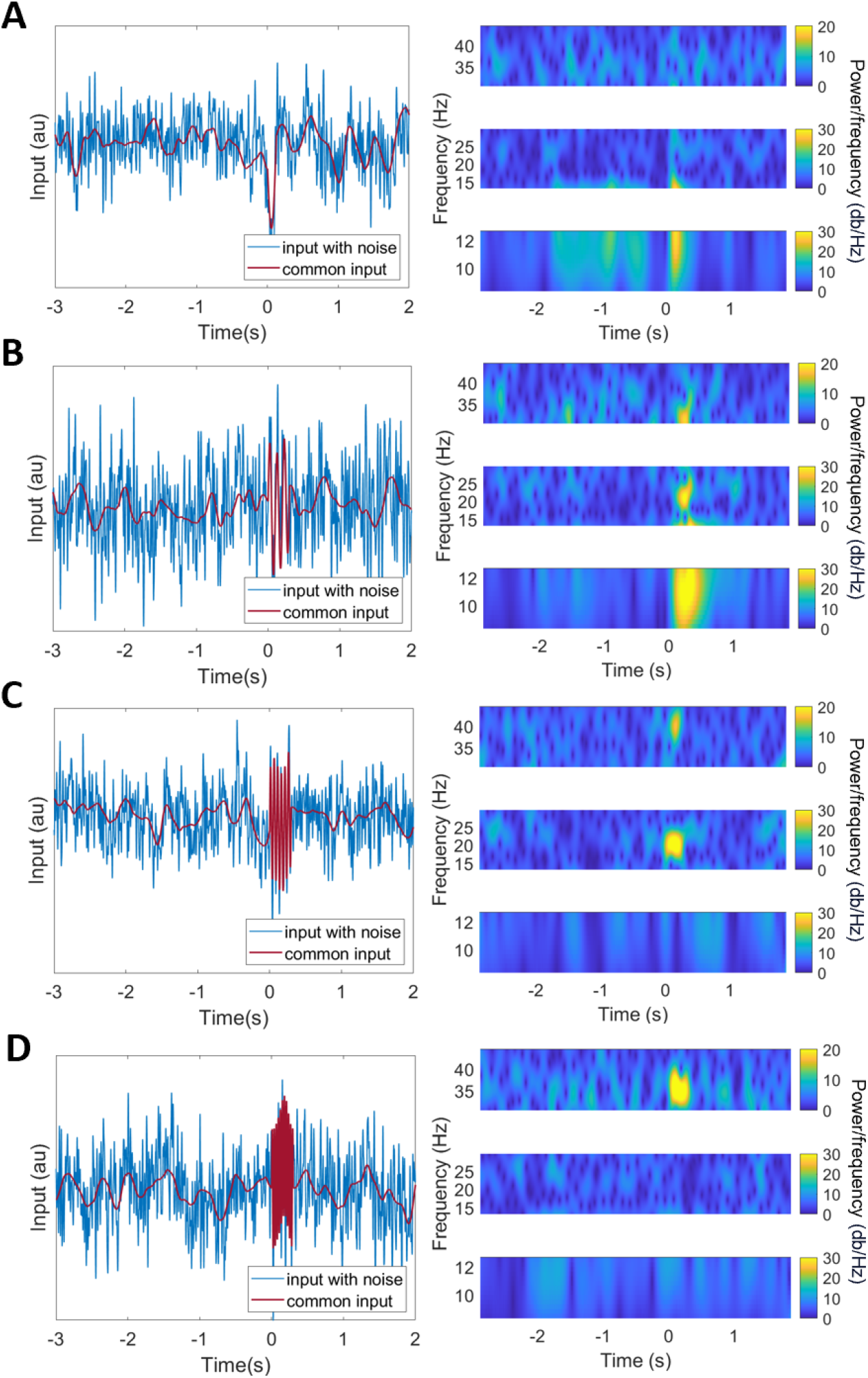
Simulations of a pool of motoneurons receiving various common inputs. A, A drop in the common input at low frequencies (red trace; left panel) causes the synchronisation of the firing activity of motoneurons, and thus increases the power at frequencies that match their average discharge rate. The spectrogram of the cumulative spike train (CST) is shown separately on the right for alpha (8-12 Hz), beta (13-30 Hz) and gamma (30-45 Hz) bands. B, Low frequency common input with an additional 300 ms oscillatory activity at 10 Hz caused an increase in power at 10, 20 and 30 Hz. C, Low frequency common input with an additional beta oscillatory input at 20Hz increased the power at 20 Hz and 40 Hz but did not affect lower frequencies. D, Low frequency common input with an additional low gamma input at 35 Hz temporarily increased the power at 35 Hz only.

In the second scenario, the pool received a short 10-Hz oscillatory input at time 0 s on top of the low frequency common drive. Figure 2B shows the common input and the spectrogram of the CST. There was an increase in power at 10 Hz, but also at the harmonics, at 20 Hz and 30 Hz, without common inputs at these frequencies. Similarly, in the third scenario, there was a 20-Hz oscillatory input to the pool (Figure 2C). Besides an increase in power at 20 Hz, the spectrogram also showed an increase in power at 40 Hz, though of lower amplitude than at 20 Hz. Finally, the last case involved a short input at 35 Hz (Figure 2D). In this scenario, the power increased at 35 Hz, without affecting the power at lower frequencies.

Overall, the simulations illustrated how a change in the low-frequency common inputs or in the high frequency inputs can synchronise the firing activity of motoneurons and how the power spectrum of the CST is affected by these events. Specifically, even a small decrease in common input at low frequencies, which may be expected with movement cancellation, would provoke synchronization of the motoneuron output at the frequencies of the average discharge rates, corresponding to the alpha band.

### Behavioural results

To confirm that participants were able to perform the task correctly, the percentage of valid trials was calculated (see Methods for criteria). On average, 96.6 ± 0.8% and 94.7 ± 3.8% of ‘GO’ and ‘NO-GO’ trials were considered valid. Subjects had an average reaction time of 275.2 ± 42.8 ms in the ‘GO’ trials.

The CoV of force was 2.8 ± 0.7% during the 4 s periods before the imperative cues in trials where force stayed between 7% and 13% MVC.

### Decomposition results

On average, 32 (range 12-45) MUs per participant were identified from EMG signals. Out of these, an average of 28 (range 10-40) MUs fired steadily (without being de-recruited and recruited) during the contraction and were considered for further analysis. The average discharge rate of these MUs was 10.5 ± 0.8 spikes/s and the CoV of the interspike intervals was 14.7 ± 1.7%.

### Changes in brain and motor unit activity

We studied changes in power in alpha, beta and gamma bands in the MU firing activity, and in the EEG during the ‘NO-GO’ trials. The grand average from all subjects showed changes in all bandwidths, while force stayed stable (Figure 3). A small drop in power was observed in the preparation period (−1 s to 0 s) and a large increase post ‘NO-GO’ cue. To explore which changes were statistically significant, repeated measures ANOVA were performed on this data.

**Figure 3:**
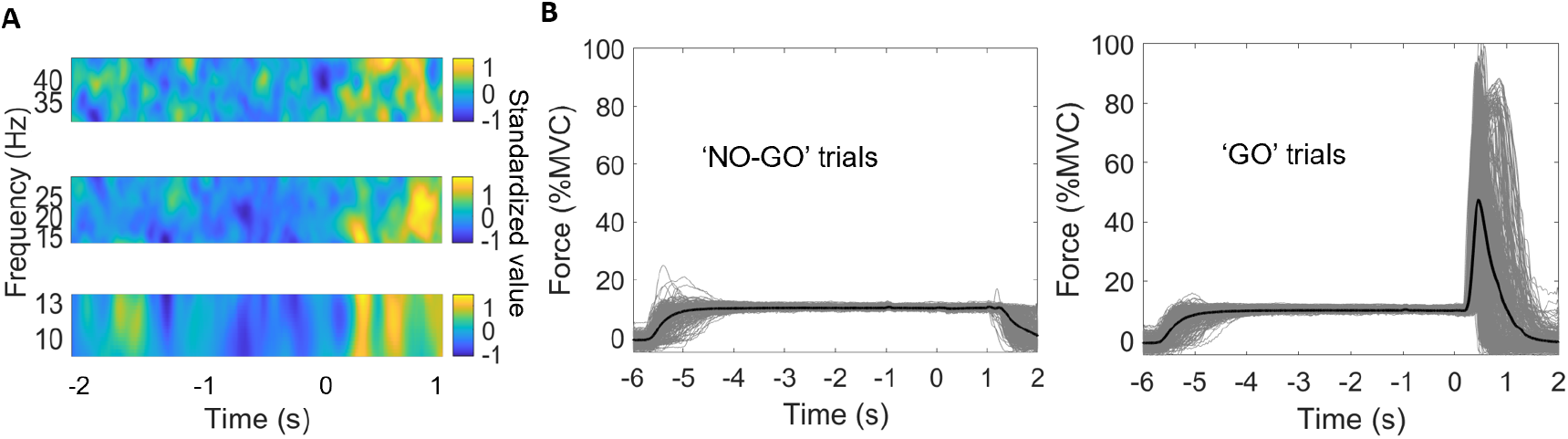
Average changes in MU activity observed while force output was kept constant. A, Standardized changes in power during the ‘NO-GO’ trials averaged from all participants. Spectrogram shows average changes in alpha, beta and gamma bands compared to the reference window (−3 s to -2 s). Values were standardized in the window shown here. B, Force traces for valid ‘NO-GO’ and ‘GO’ trials. Individual trials from all participants were plotted in grey and average in black.

Overall, at the MU level, we found an effect of time on power in alpha (F = 8.19; p = 0.004; *η*^2^ = 0.51), power in beta (F = 29.86; p < 0.001; *η*^2^ = 0.79) and power in gamma (F = 9.69; p = 0.002; *η*^2^ = 0.55). At the brain level, there was a significant effect of time on power in beta (F = 17.28; p < 0.001, *η*^2^ = 0.68) and gamma (F = 18.60; p < 0.001; *η*^2^ = 0.70), but not alpha (F = 0.51; p = 0.61; *η*^2^ = 0.06). There was no significant effect of time on the average discharge rate of decomposed units (F = 1.28; p = 0.304; *η*^2^ = 0.14).

In the *preparation* period compared to *baseline*, we did not observe any significant changes in high frequency oscillatory activity at the brain (beta: p = 0.699; gamma: p = 1.000) or muscle recordings (beta: p = 0.159; gamma: p = 0.265) (Figures 4 & 5).

**Figure 4:**
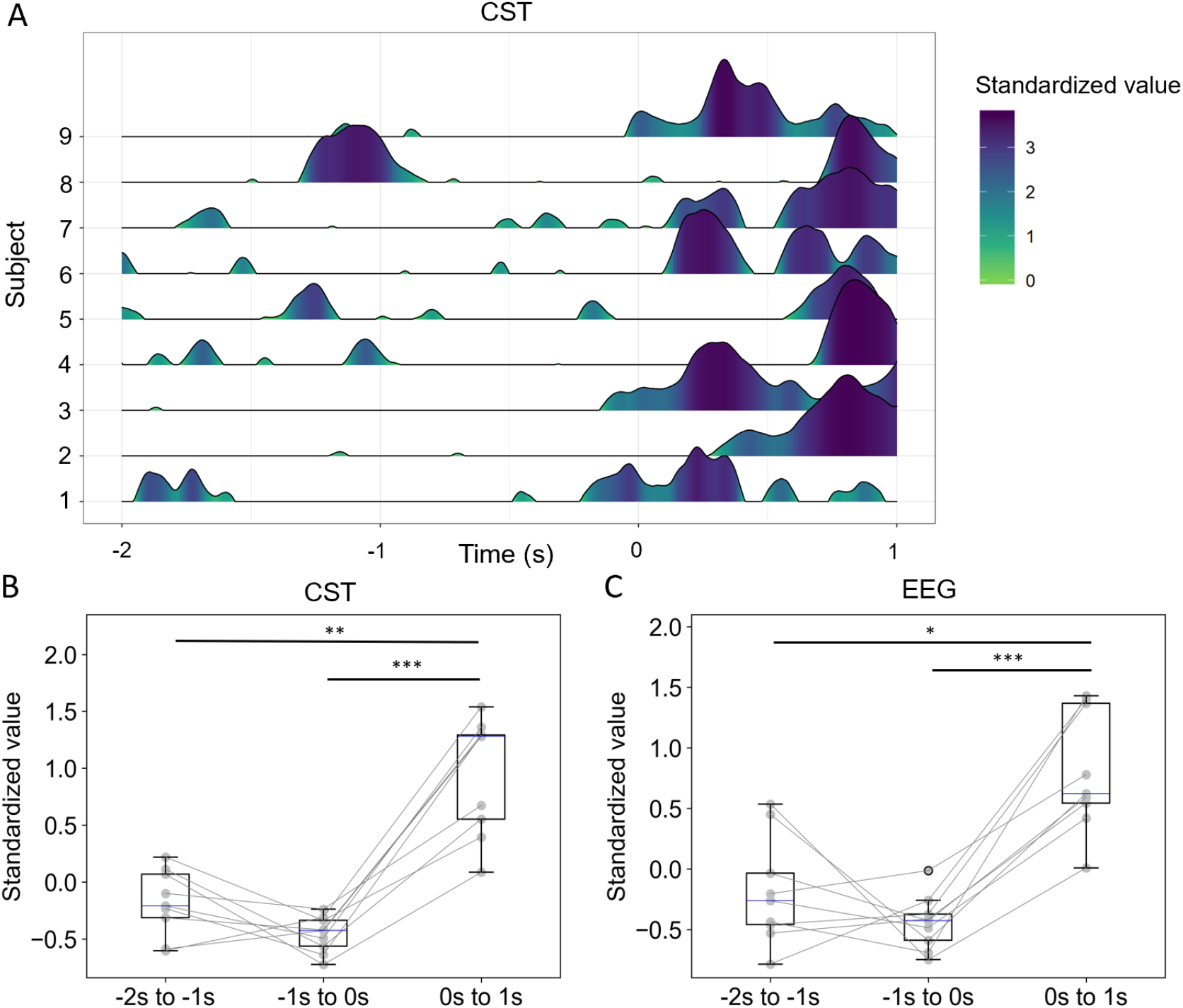
Changes in beta (13-30 Hz) levels at MU and brain levels. A, Evolution of beta power in ‘NO-GO’ trials at muscle level. Each row represents data from one subject. Values were standardized in the window -2 s to 1 s. Only positive values are plotted for clarity. B, Changes in beta band MU activity in three windows relative to the ‘NO-GO’ cue. Each point and line represent one subject. There is a significant increase in the last window compared to the first two. C, Changes in beta observed in EEG recordings. Beta levels are significantly increased post ‘NO-GO’ cue. *p < 0.5; **p < 0.01; ***p < 0.001

**Figure 5:**
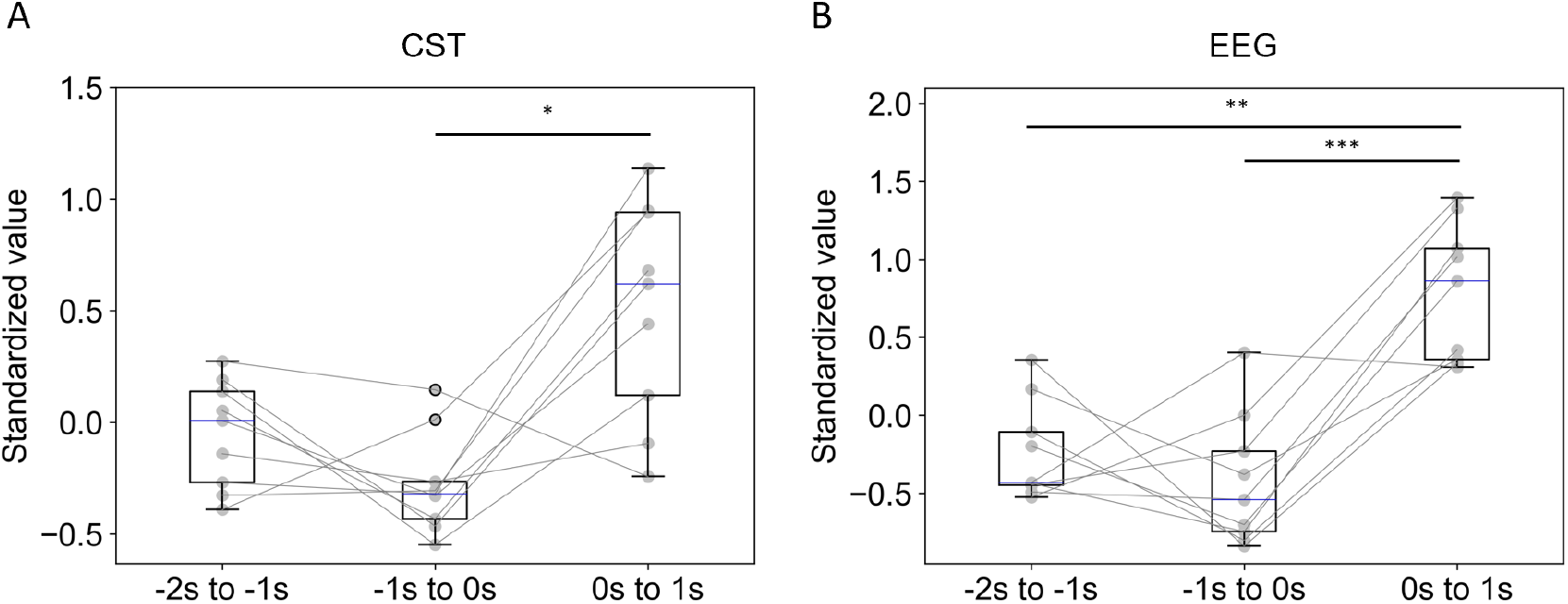
Changes in gamma (30-45 Hz) levels at MU and brain levels. A, Average gamma level in MU activity in the three windows. Each point and line represent data from one subject. There is a significant increase in the preparation cancellation period compared to the previous window. B, Gamma levels observed in EEG recordings. There is a significant increase in the last window. *p < 0.5; **p < 0.01; ***p < 0.001

In the *cancellation* period, we observed a brief drop in the average DR (effect observed in six out of the nine subjects), that caused a synchronization in motoneuron firings. This observation was similar to the first simulation scenario described above, where a drop in common input caused a drop in DR, followed by synchronization of units. At approximately the same time of the drop in DR in the physiological data, there was also an increase in the average power of lower frequencies (8-12 Hz) in the MU activity, which was not present in the EEG activity. Due to this change, there was a significant increase in the power within 8-12 Hz during the *cancellation* window compared to the *preparation* period (p = 0.003). This observation was fully predicted by the simulations.

We also found a difference in average beta levels in the MU activity, with the *cancellation* interval showing a significant increase in beta power relative to *baseline* (p = 0.008) and *preparation* (p < 0.001) (Figure 4B). The timing of these rebounds was approximately aligned across subjects (Figure 4A). To explore whether these changes had a cortical origin, the same analysis was performed on the EEG recordings (Figure 4C). EEG data showed similar patterns to what we saw in the MU activity, with an increase in beta levels after the ‘NO-GO’ cue. Beta power was significantly higher during *cancellation* (0 s to 1 s) than during *baseline* (−2 s to -1 s) (p = 0.028) and *preparation* (−1 s to 0 s) (p < 0.001) in all subjects (Figure 4C). The peak amplitude of the forward coherence (EEG *→* MU) was also measured in the different windows considered. Four out of nine subjects had significant coherence in beta during *baseline* (0.19 ± 0.14), three subjects during *preparation* (0.19 ± 0.11) and seven subjects in the last window (0.18 ± 0.12). Four subjects had non-significant coherence during *preparation*, that became significant post ‘NO-GO’.

Similar changes to the ones observed in the beta band were found in the low gamma band (30-45 Hz) activity (Figure 5).There was a significant increase after the ‘NO-GO’ cue compared to preparation in MU activity (p = 0.011), with the change not being significant compared to baseline (p = 0.113) (Figure 5A). In the EEG, the power in gamma was significantly increased in the last window compared to both first and second windows (p = 0.007 and p < 0.001) (Figure 5B). Three out of nine subjects had significant forward coherence in the low gamma band during baseline (0.05 ± 0.02), four subjects during preparation (0.05 ± 0.01) and five subjects in the last window (0.07 ± 0.01).

Overall, we found a significant increase in beta and low gamma at both muscle and brain level, while the changes in low frequencies (< 10 Hz) were only present in the MU activity.

## Discussion

We studied the transmission of cortical inputs to spinal circuits during movement preparation and cancellation. We showed that the changes in high-frequency oscillatory activity in the beta band previously reported in brain recordings during movement cancellation is also present in the MU activity. This study thus provides evidence for the possibility of extracting changes in brain states from the signals recorded at the periphery of the nervous system.

The initial part of the study explored, through simulated data, how changes in common inputs to motoneurons can modulate their firing activity during two scenarios: i) a significant change in the low frequency common input that directly affects the motoneuron discharge rate, or ii) an increase in higher-frequency oscillatory common inputs that does not impact the motoneuron discharge rate. Both a drop in low frequency common drive and an added oscillatory input affected the power of the CST at various frequencies. More specifically, in the first scenario, the increase in power was most evident at the frequency of the DR values, while in the second scenario, the effect corresponded to the frequency of the oscillation and its harmonics. The results of the simulations were used to interpret the experimental results where we do not have direct access to the inputs to spinal motoneurons.

Through physiological recordings, we then studied how spinal motoneuron population activity is modulated during movement preparation and cancellation, while force is held constant. Changes in cortical activity during these states have been reported in previous studies (Schoffelen et al., 2005; Schoffelen et al., 2011; Wessel, 2020), but here we focused on the transmission of this activity to spinal motoneurons. The results add to our knowledge on what type of information, and in which conditions it is transmitted from the brain to the muscles. They also corroborate the concept that the information travelling from the cortex during various motor stages does not necessarily have a direct impact on behaviour.

In the experimental data, we observed a significant increase in beta power during the *cancellation* period, which matched changes recorded at the brain level. Human studies looking at changes in the beta oscillatory activity using EEG recordings have previously reported increases in this frequency band after movement cancellation (Alegre et al., 2004; Solis-Escalante et al., 2012; Wessel, 2020). In a stop-signal task, where participants had to rapidly cancel a movement they were previously prompted to do, beta bursting increased after successful cancellation (Wessel, 2020). In a similar ‘GO’/’NO-GO’ paradigm as the one used here, Alegre et al. (2004) reported beta synchronization in the fronto-central brain areas post ‘NO-GO’. Our results are in line with these previous findings, even though there is one crucial difference between the tasks of most previous works and ours. In previous studies, subjects were relaxed while preparing for movement, while in the current work, participants were required to hold a stable contraction, so MU activity could be extracted in the baseline condition. Another study used a similar paradigm where local field potentials were recorded from monkeys while they were doing a lever depression task in a ‘GO’/’NO-GO’ paradigm (Zhang et al., 2008). The monkeys had to initiate a lever depression and react to visual patterns by either releasing the lever (‘GO’) or maintaining the motor output. In the ‘NO-GO’ trials, the authors reported a marked beta rebound after a desynchronization that appeared in all types of trials (Zhang et al., 2008).

Overall, previous results clearly indicate that movement cancellation is characterized by an increase in beta power cortically, independent of the initial state (rest or stable contraction). Critically, here we show that this change in beta activity is transmitted to the spinal motor neurons. The coherence in seven out of the nine participants during the *cancellation* period further supports the cortical origin of these changes. The lack of significant coherence in the other two participants might results from the computation of coherence over a 1 s window, while beta activity might be transmitted in shorter bursts (Echeverria-Altuna et al., 2022; Little et al., 2019; Wessel, 2020). During the *preparation* period we observed a small decrease in beta power in some of the participant that was not significant. This would be in line with the idea of motor system disinhibition while preparing a movement (Engel and Fries, 2010).

Power in lower frequencies (8-13 Hz) also increased in the last window at the MU level, but not at the brain level. This suggests that this increase in power was not due to the modulation of cortical descending oscillatory inputs to muscles. These frequencies match the average discharge rate of the identified MUs. Thus, this increase in power might be due to the synchronisation of the firing activity of MUs through a transient change in common inputs at lower frequencies. The short drop in discharge rate also points towards a brief drop in common drive to the MUs. This phenomenon is illustrated in one of the simulation cases presented at the beginning of this work in Figure 2. Although a non-cortical oscillatory input could also increase power in alpha, the drop in DR we observed suggests that the first scenario is more likely. We also observed an increase in power for the low gamma band (30-45 Hz) during the *cancellation* period at both brain and muscle levels. The level of forward coherence (from brain to muscles) was significant in half of the participants during this window. Previous work reported an increase in coherence over the gamma band during movement preparation (Schoffelen et al., 2011), which was not observed here. However, one key difference between the paradigms used in studies that reported gamma activity in preparatory periods and our work is the level of unpredictability of when the cue was presented (Schoffelen et al., 2005; Schoffelen et al., 2011). We used a fixed interval between the warning and imperative cues, while in previous studies, the timing of the cue subjects had to respond to was varied. This might suggest that gamma, in the context of preparation, could be a sign of unpredictability. We however, observed a change in gamma (30-45 Hz) post ‘NO-GO’ cue in both EEG and MU activity. One concern is that these peaks in the spectrogram of CST could be partly produced as harmonics of other oscillatory synchronizations. This phenomenon was observed in the simulations of oscillatory common inputs at 10 and 20 Hz (Figure 2). Although it remains possible that strong beta synchronisation observed during the cancellation period could have affected the level of power in the gamma band, note that we also found significant forward coherence during this window in five out of nine participants. This suggests that there is also a transmission of these descending oscillatory inputs to muscles at these frequencies, but the effects are not as strong as in the beta band. In addition, even though we identified on average 28 steadily firing MUs per participant, some of them had significantly fewer units. Thus, the estimation of power in higher frequencies, especially in the gamma band, could have been affected by their limited sampling by the CST of the identified motor units.

Overall, we observed significant changes in the firing activity of spinal motoneuron during movement cancellation, while the force output was held constant. Two mechanisms might explain the results observed here: synchronisation through a drop in input and high-frequency cortical oscillatory rhythms leaking to the motoneurons. The consistency of changes observed in the beta band at brain and muscle levels provide evidence of the effective transmission of cortical oscillatory inputs to the MUs during changes in brain state, but stable contractions. Moreover, the average discharge rate did not change during the three periods (*baseline, preparation, cancellation*), further supporting the idea that the significant changes in high frequency modulation did not have a functional effect on force production and behaviour. This further raises the question of the functional relevance of these high frequency rhythms. One hypothesis is that beta oscillatory activity has a role in transmitting state-related information by feedback to the brain (Baker, 2007; Witham et al., 2011), which is not in contradiction with our results, though such claim cannot be supported by our data.

Finally, the transmission of cortical oscillations substantiated in this study supports the idea that muscle readings effectively provide information on brain activity. The oscillations identified from MUs could be used to estimate the corresponding supra-spinal oscillations. The motoneuron, as the final common pathway of the neuromuscular system, receives a variety of inputs from the full system, which could allow researchers to partly infer the activity of other regions of the nervous system, including brain activity that does not directly drive muscle force.

## Acknowledgements

The authors are grateful to Agnese Grison for her help with the intramuscular electromyography data acquisition.

## Author Contributions

BZ: Conceptualization, Methodology, Investigation, Analysis, Writing – original draft, review & editing; SA: Investigation, Writing - review & editing; JI: Conceptualization, Methodology, Analysis, Writing – review & editing; DF: Conceptualization, Resources, Writing – review & editing

